# Integrated flexible DNA methylation-chromatin segmentation modeling enhances epigenomic state annotation

**DOI:** 10.1101/2025.07.25.666820

**Authors:** Nihit Aggarwal, Johanna Elena Schmitz, Lukas Laufer, Sven Rahmann, Jörn Walter, Abdulrahman Salhab

## Abstract

DNA methylation and histone modifications together shape the cell-type-specific epigenomic landscape. To enable a more comprehensive genome-wide annotation, we developed EpiSegMixMeth (ESMM), the first truly integrative segmentation model combining chromatin marks and DNA methylation. ESMM extends hidden Markov models with flexible read count distributions and state duration modeling. Applied to 154 high-quality human epigenomes from the IHEC EpiAtlas, ESMM substantially improves the annotation of broad heterochromatic regions-covering over 60% of the genome, that are frequently missed by chromatin-only models. Additionally, it precisely defines the boundaries of narrow regulatory elements and resolves local chromatin state transitions during cell differentiation. Notably, we demonstrate that DNA methylation can substitute for missing repressive histone marks in segmentation, ensuring robust annotation across diverse cell types. In memory B-cell development, ESMM reveals fine-scale chromatin state shifts that align with 3D genome architecture changes. Our results highlight the power of integrating DNA methylation into genome segmentation and provide a valuable resource for dissecting cell-type-specific epigenomic regulation.

## Introduction

The epigenomic landscape of the genome is comprised by cell-type-specific modifications. A complex combination of DNA methylation and post-translational histone modifications establishes a chromatin *grammar* linked to the regulation of gene expression and the organization of the genome into distinct functional sub-compartments in the nucleus [1, 2, 3]. Over the last decade a number of comprehensive cell-type-specific epigenomes have been collected by national and international consortia. These are currently assembled in a first comprehensive human epigenome cell atlas by the international human epigenome consortium IHEC [4, 5]. This data is essential to understand the cell type and cell state specific programs executed from a unique genome. Currently, a comprehensive epigenome is defined by DNA methylation, RNA-seq and (a minimum of) six core post-translational histone modifications that establish the cell-type-specific genome programs [1, 5].

Epigenetic modifications operate at different scales ranging from individual base pairs (DNA methylation) to nucleosomes up to large chromatin domains. There is a significant overlap and crosstalk between these marks [6, 7]. For instance, high cytosine methylation correlates with H3K9 methylation [8, 9], while it anticorrelates with the regulatory mark H3K4me3 and in regulatory regions additionally with the polycomb-associated repressive mark H3K27me3 [10, 11, 12, 13, 14].

Current segmentation and genome annotation (SAGA) algorithms computationally model these complex relationships [15] as a multidimensional linear sequence of epigenetic states and link them to functionally annotated genomic regions, such as promoters, enhancers and transcriptional states [16, 17, 18]. In other approaches, the three-dimensional organization of genomes derived from Hi-C data has been transformed into linear segments (compartments) and linked to chromatin states to better understand the role of the spatial nuclear (chromatin) context, generating a three-dimensional view on the functional organization of gene regulation and chromatin crosstalk [19, 20, 21]. All widely used segmentation tools [15] either exclusively use histone marks as input, such as ChromHMM [18], Segway [22, 23], EpiCSeg [17], and EpiSegMix [16], or only rely on cytosine methylation signals, such as MethylSeekR [24], Methpipe [25], and MMSeekR [26]. As a result, the biological state annotations differ between these two classes of segmentation tools and are rarely combined. For instance, in the context of chromatin segmentation, terms such as promoter, enhancer, or transcribed regions partially overlap and correspond to DNA methylation states that are categorized as unmethylated, lowly methylated, or highly methylated domains, respectively. Moreover, CpG positions that contain the DNA methylation information are not equally distributed across the genome, generating regions with low or no information content. The generation of an integrative model that effectively incorporates both classes of epigenetic signals and models their interactions would thus offer a more comprehensive and interdependent view on the epigenomic landscape.

As a basis for such an integrated segmentation tool, we use the basic framework of EpiSegMix (ESM) [16], a recently developed segmentation tool which incorporates new features to address some of the limitations of existing tools. First, ESM has a broader flexibility to model the distribution of epigenetic marks, by not only supporting the conventional Poisson or Negative Binomial distribution, but a wide range of discrete probability distributions with varying flexibility to model variance and skewness of the histone count data. As a consequence, it allows us to better capture the distinct distributional properties of histone mark counts, such as the often observed overdispersion of H3K27me3 or H3K4me1 counts. Second, built-in flexible duration modeling efficiently captures short-duration (nucleosomal) and long-duration (domain) states simultaneously in a single model.

We here present an extended version of ESM, which we name EpiSegMixMeth (ESMM). ESMM is a comprehensive epigenomic segmentation tool addressing the challenges for an integrated probabilistic model for a truely combined and integrated use of genome wide chromatin data and DNA methylation data. We apply ESMM to a large and uniquely processed full epigenome dataset provided by the IHEC Human Epigenome Atlas [4, 5]. To our knowledge, ESMM is the first tool to support such a read count based simultaneous integration of different epigenetic modalities. We show that ESMM enhances the cell-type-specific classification of epigenomes particularly in large, previously not well captured and annotated heterochromatic regions. As an example, we show how ESMM data can be used to follow epigenomic state transition in developing cells revealing new interpretation links to three-dimensional Hi-C data.

## Results

### ESMM: An integrative segmentation tool for histone and methylation data

To generate an integrated segmentation model we combined approaches developed for DNA methylation segmentation with our recently developed chromatin based tool EpiSegMix (ESM) [16]. In short, we constructed a flexible extended Hidden Markov state duration model that incorporates both histone modifications and DNA methylation sequencing data on a read count-based level (see Methods for further explanation). Epigenetic signals were captured in non-overlapping windows of 200 bp. Histone mark counts were obtained by taking the number of reads mapping to a window, while for DNA methylation both CpG coverage and methylated CpGs falling into a window were considered. In addition, ESMM incorporates duration modeling to account for heterogeneous segment length distributions throughout the genome [16].

To evaluate the performance and robustness of ESMM, we applied it to 154 complete, non-imputed epigenomes from healthy human cell or tissues, uniformly processed and obtained from the IHEC EpiAtlas [4]. Each complete epigenome consists of whole genome bisulfite sequencing (WGBS), RNA sequencing (RNA-Seq) and six core histone marks: H3K4me3, H3K27ac, H3K4me1 (narrow marks), H3K36me3 (transcription mark), and broad repressive marks H3K27me3 and H3K9me3. The sample set spans multiple tissues and primary cells grouped into 19 distinct cell types (see Figure 1A), including a large set of immune cells (see Additional Figure S1). While some variability in signal quality and quantity was observed (see Additional Figures S2A-C), the overall data quality was deemed sufficient for integrative modeling.

**Figure 1:**
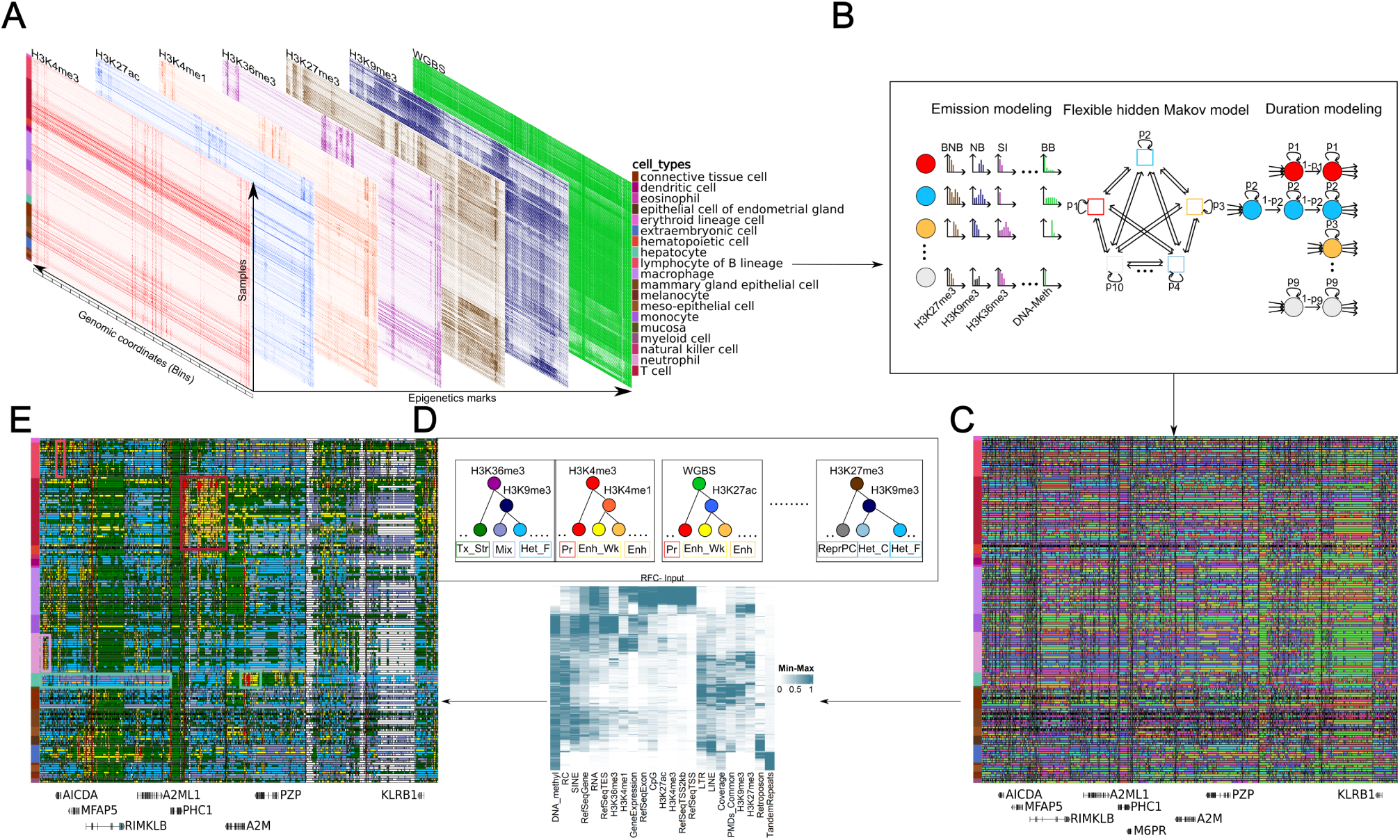
Schematic overview of the joint-model and annotation strategy. (A) Histone modifications and DNA methylation signals across 154 human epigenomes are used as input for the segmentation model. Samples span a wide range of primary tissues and cell types. Segmentation was performed using the human hg38 reference genome and a bin size of 200 bp. The region shown corresponds to AICDA gene locus (chr12:8550000-9650000). (B) The EpiSegMixMeth (ESMM) model fits an HMM with mixture and duration modeling, trained independently for each sample. (C) Chromatin states across 154 epigenomes in AICDA locus using 10-state EpiSegMixMeth (ESMM) model. State labels are assigned arbitrarily per sample. (D) A random forest classifier was applied to each sample to map the numerical state labels to unified, biologically meaningful chromatin states. (E) AICDA locus for the same samples post uniform labeling, providing clear cell-type-specific states as well as agreement between samples of the same cell and tissue type. Pink, red, and light purple boxes surround active regulatory states for B-, T-cells and macrophages, respectively. Turquoise box encapsulate liver-specific repressive states (left) and liver-specific active regulatory states (right).

Each of the 154 samples was modeled independently. Based on empirical testing with different numbers of states, we found that a 10-state model provided the most stable and biologically interpretable segmentation across all samples.

### Generation of unified ESMM state annotations across epigenomes

As a consequence of the individual training for each sample the ten hidden states are arbitrarily ordered in each trained model. Hence, to enable direct comparison of chromatin states across all 154 samples (see Figure 1C), we implemented a unified state classification method. We employed a random forest classifier to identify and label the states across all samples (see Figure 1D and Methods section for more details). This classifier used 23 features, including the average signal and overlap enrichment across six histone marks, DNA methylation, genomic coverage, gene expression, genes, transcription start site (TSS), transciption end site (TES), common partially methylated domains (PMDs) [27], and repeat elements [28].

The unified labeling, followed by grouping cell types, revealed distinct cell-type-specific state emission patterns (Figure 1D). For example, Figure 1E highlights a genomic region containing genes active in lymphoid cells where we observed strong enrichment of regulatory states (yellow and orange states in the pink, red, and light purple boxes). These include B-cell-specific expression of AICDA gene (see Additional Figure S4A), T-cell specific expression of KLRB1 [29], and macrophages-specific activity of CLEC4E [30], each associated with regulatory chromatin states.

Importantly, these cell-type-specific patterns extended beyond activation signals. We also observed widespread cell-type-specific repressive states (light blue and purple states in turquoise box), often characterized by an absence of gene expression. Notable examples include clear repressive chromatin configurations in liver cells at immune-cell-specific loci such as CLEC4E, AICDA, and others (see Additional Figure S4B).

In summary, the flexible segmentation provided by ESMM enabled the construction of a comprehensive, high-resolution map of chromatin states across diverse cell types. It successfully captured both activating and repressive patterns, offering key insights into cell-type-specific epigenomic programs.

### Integrative epigenome segmentation recovers large domains missed by histone-only models

A comparative analysis of the genome-wide state enrichment across 154 samples reveals that ESMM performs well not only in gene-rich regions and regulatory spaces, but also in large parts of the genome which ChromHMM, a histone-only-based segmentation model, assigns the *quiescent* or *no signal* state (see Figure 2 A-B). ESMM segments a large proportion of these previously unannotated regions to various functional heterochromatic states (see Additional Figure S4B as an example region). We notice that ChromHMM defines exclusive states which exhibit a lower certainty for all labels during classification. We annotate these three ChromHMM-exclusive states (marked with asterisk in Figure 2C) as transcribed (Tx), weak promoter (Pr Wk), and repeat (Rep) states. In comparison, ESM (see supplementary Figure S5) and ESMM define a specific (mixed) state not covered by ChromHMM. In total, the comparison between ESMM and ChromHMM comprises 13 distinct functional states. The exclusive ChromHMM states Tx, Pr Wk and, Rep are submerged in other states of ESMM leading to clusters of regulatory domains with slightly variable sample counts but highly similar patterns. Despite of this seemingly lower granularity, ESMM is best in differentiating between low and high transcriptional active genes (see below). The high transcribed state is marked by high H3K36me3 and high DNA methylation levels. This Tx Str state covers relatively larger domains (see Figure 2A). ESMM utilizes overlapping features of chromatin and DNA methylation and detects a significant co-occurrence of features and terminologies provided by chromatin and DNA methylation (e.g. unmethylated regions (UMRs) and lowly methylated regions (LMRs)) based segmentations. We noticed that the Tx state defined by ChromHMM is inconsistent between samples, which is reflected in the genomic coverage in Figure 2B and the Tx Wk state shows relatively low occupancy.

**Figure 2:**
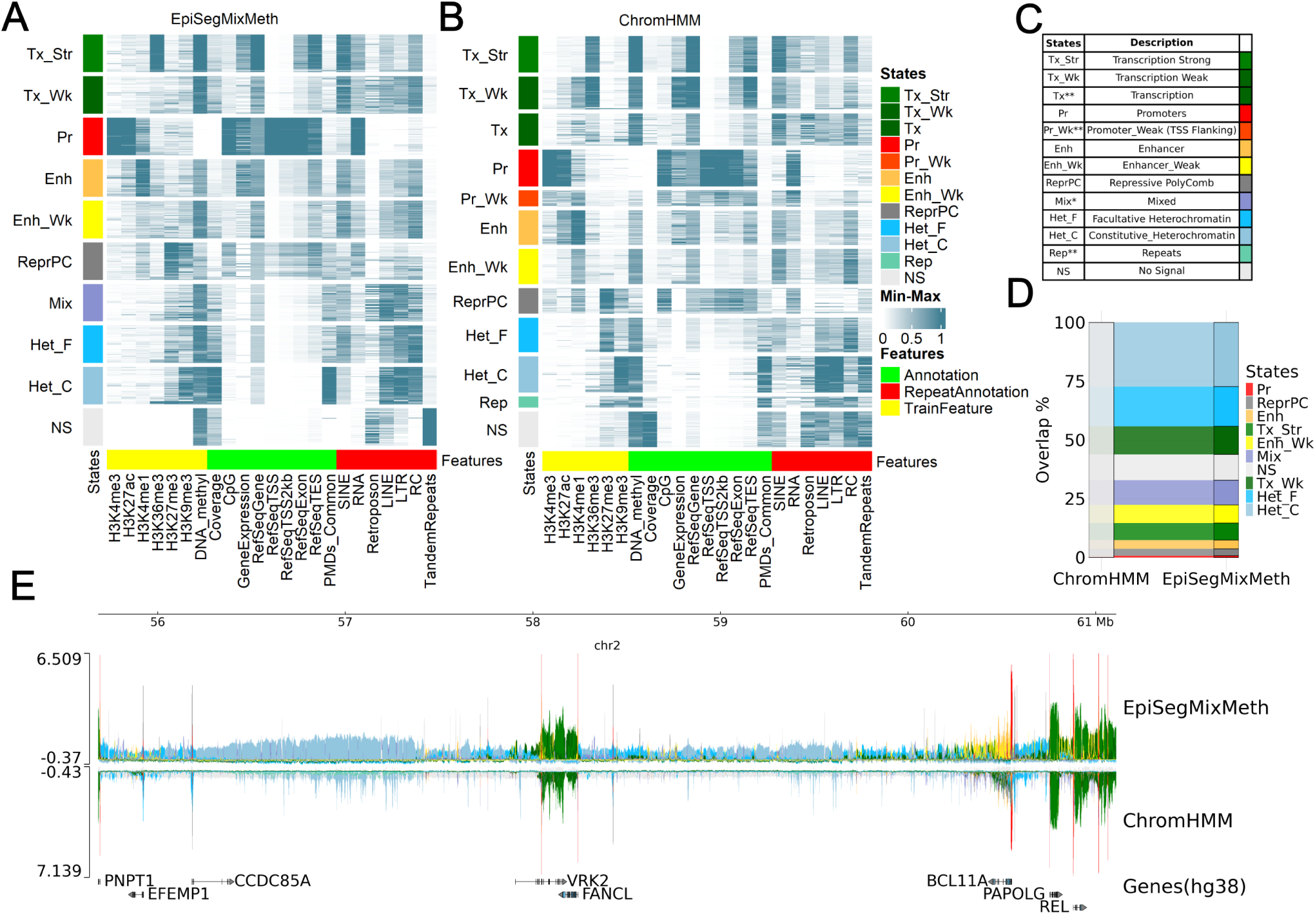
Epigenomic-wide state characterization of 154 samples. (A) ESMM states defined using six core histone marks and DNA methylation annotated with biological functions. Labels are assigned based on the random forest classifier. Features on the x-axis are grouped into *train* (yellow), which are used in the model; *annotation* (green); and *repeat annotation* (red). (B) ChromHMM states from models trained only on the six core histone marks, assigned labels same as A. Note that large part of the genome is assigned to No Signal state (NS). (C) States with their biological associated function and color code. ChromHMM-specific states are denoted with asterisk **, while ESMM-specific state is denoted with *. (D) Overlap between ESMM states and ChromHMM NS state, showing re-classification of ChromHMM-NS states to various functional biological states in ESMM. (E) epilogos: a multi-biosample functional genomic annotations view around a large heterochromatic domain locus (chr2:43832201-62396766) for ESMM and ChromHMM. Y-axis shows the S1 score of epilogos for ESMM whereas the inverted y-axis for ChromHMM. S1 quantifies deviation of chromatin state frequency from the genome-wide average; higher scores indicate atypical/important patterns across biosamples.

The promoter state (Pr) is consistently captured in all models, in line with a very low methylation in CpG dense promoter regions (UMRs) and the highest enrichment of the signature promoter marks H3K4me3 and H3K27ac compared to all other states. Both, ESMM and ChromHMM, define two enhancer states (Enh and Enh Wk) with apparent differences in H3K4me1 and DNA methylation levels. Weak enhancers show high DNA methylation levels and a reduced/depleted H3K4me1 enrichment. Overall, we observe a very good agreement between ChromHMM and ESMM euchromatin-specific states, including transcription and regulatory states, which cover a relatively small part of the genome together.

A major difference between ESMM and ChromHMM emerges as ESMM provides a much broader coverage and clearer distinction of heterochromatic states. The repressive polycomb (ReprPC) state, is marked by high H3K27me3 and an intermediate level of DNA methylation and CpG density (see Figure 2A). ESMM features a novel mix state of comprising a combination of the repressive marks H3K27me3 and H3K9me3, an enrichment of repeat elements and a moderate DNA methylation. The facultative heterochromatin state shows a relatively lower enrichment for H3K27me3, reduced DNA methylation including some common (shared) PMDs and favors rolling circle amplification (RC). While ChromHMM also identifies these states, the enrichment is less pronounced and the genomic distribution and coverage appear more unstable. The most repressed state, constitutive heterochromatin (Het C), shows the highest enrichment of H3K9me3, a constant high CpG-methylation signal and an enrichment of long terminal repeats (LTRs). Constitutive heterochromatin shows an almost exclusive enrichment of common (shared) PMDs (defined in [27]) pointing towards a substantial overlap between ESMM and MethylSeekR, a methylation-based, segmentation approaches.

In summary, ESMM enhances the quality and quantity of state assignments throughout the genome. The proportion of the genome classified as *no signal* state is reduced from up to 70% by ChromHMM to about 10% (see Figure 2D and as an example supplement Fig 8 A/B). ESMM not only enhances the overall calling of heterochromatic segments (Figure 2E), it also differentiates no signal states into either constitutive or facultative heterochromatin or a transitional mix state. Overall, ESMM provides an enhanced and robust genome wide classification of heterochromatin not only in large heterochromatic domains but also in smaller intergenic regions as can been seen of generally higher epilogo scores associated with a facultative heterochromatin (and mix) state (see an example Figure 2E).

### DNA methylation supports robust calling of eu- and heterochromatic states

We conducted several analyses to better understand and assess the contribution of DNA methylation into ESMM. First, we evaluated gene expression prediction accuracy (measured by R-square) based on functional chromatin states inferred from three different models: ChromHMM, ESM, and the integrative ESMM model. As shown in Figure 3A, ESMM showed a slight improvement over ESM and a more substantial performance gain compared to ChromHMM. Next, we investigated to what extent DNA methylation substitutes for missing heterochromatic marks. For this, we used a dataset of 40 high quality T- and B-cell immune cell samples and compared model performance under two settings: (i) complete input with all six histone marks, and (ii) het-reduced input missing H3K27me3 and H3K9me3 (the heterochromatin marks). We compared the state assignments from the reduced models of ChromHMM, ESM, and ESMM to those from their respective complete models using the Jaccard similarity index and genome-wide coverage.

**Figure 3:**
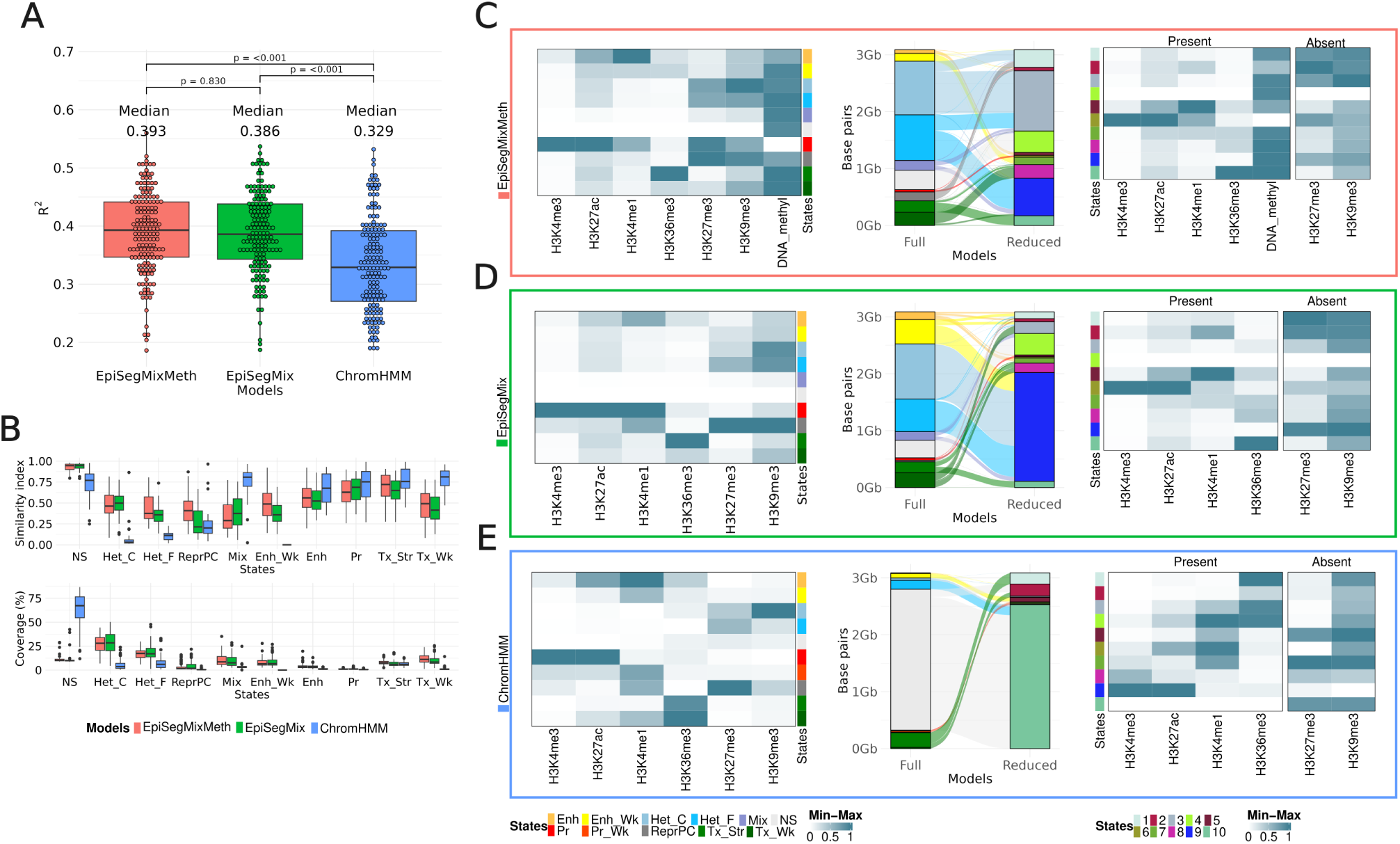
Contribution of DNA methylation to the mixture model segmentation. (A) Dotplot and boxplot measuring predicted gene expression (R-square score) accuracy using functional states defined by three different models. (B) The upper panel shows genome-wide similarity. The Jaccard index measured between the same states, defined using the full and reduced models for 40 B- and T-cell samples. The complete model consists of all training features used in each respective model, i.e., six core histone marks (H3K4me3, H3K27ac, H3K4me1, H3K27me3, and H3K9me3) and DNA methylation for EpiSegMixMeth whereas six core histone marks for EpiSegMix and ChromHMM. The reduced model removes the broad repressive marks H3K27me and H3K9me3. The similarity index represents the fraction of each state that can be captured efficiently even in the reduced model compared to the full model. The lower panel shows the genomic coverage of the corresponding state in a full model across all samples. (C) left: Naive B-cell heatmap showing enrichment of the training features for a complete model with 10 functional states defined by EpiSegMixMeth, middle: Sankey plots displaying overlap in basepairs between states defined by the full and reduced models, right: Naive B-cell heatmap for reduced model states defined by EpiSegMixMeth with additional enrichment of absent marks in the model marked as Absent. (D) and (E) are same as (C) for defined states between full and reduced model using EpiSegMix and ChromHMM, respectively.

Despite the absence of heterochromatin marks, ESMM accurately recovered heterochromatic and no-signal states (see Figure 3B). It also maintained high fidelity in detecting weak enhancers and transcriptional states, suggesting that DNA methylation provides compensatory information, particularly in regions associated with heterochromatin and euchromatin.

Notably, for most chromatin states, ESMM in the reduced setting exhibited the highest overlap with the full model among all methods tested. Each state in the reduced ESMM model corresponded clearly to a specific state in the complete model – a consistency not observed with ESM or ChromHMM (Figure 3C-E) (shown for a representative naive B-cell sample).

Moreover, the emission matrix of the reduced model of ESMM displays a similar enrichment pattern as the complete model, even for the marks excluded from training (H3K9me3 and H3K27me3) (Figure 3C). This illustrates the potential of DNA methylation to compensate for absent heterochromatic marks and accurately recover key repressive chromatin states, including repressive polycomb (ReprPC), facultative heterochromatin (Het F), and constitutive heterochromatin (Het C).

Although ESM without DNA methylation can already recapture the active state to some extent, it failed to differentiate between the different classes of closed chromatin states. Instead, it assigns all of them, along with the weak enhancer state, to one state, as shown in the sankeyplot in Figure 3D. ChromHMM, in the reduced model, categorizes –depending on the cell type– up to 70% of the genome into one state (*no signal*), overlapping with the *no signal* states, facultative and constitutive heterochromatin states in the full model.

Having shown that DNA methylation can be a proxy for the absence of other chromatin marks, we next examined whether DNA methylation improves segmentation stability, e.g., by compensating for an inconsistent or lower quality of chromatin immunoprecipitation followed by sequencing (ChIP-Seq) signals (Additional Figure S2). To demonstrate this, we analyzed segmentation results in relation to the variation in the quality of H3K27me3. We compared the individual sample state performance against robustness score using the Jaccard similarity index and genomic coverage for states affected by this mark in a set of luminal epithelial cells from mammary gland samples.

We observe that the states defined by ESMM are more stable and show less variability against the quality changes of H3K27me3 as compared to a segmentation by ESM or ChromHMM (Additional Figure S6A). The genomic coverage of ESM, defining facultative heterochromatin, ranges from 1% to 50% for the low-quality H3K27me3 samples (below 30 Jenson-Shannon Distance (JSD) score), while ESMM produces a more stable annotation of these states (7% − 30%). We observed that the overall variance for the proportion of Het F state varied primarily based on the JSD score and was more prevalent in ESM than in ESMM. As an example, we highlight (see Additional Figure S6A) a set of luminal epithelial cells of the mammary gland. In our initial QC we noticed that these samples fall into sets of low, mid, and high quality levels across eight replicates. Interestingly, the proportions of the heterochromatin state were robust across all replicates for the ESMM mixture model, while ESM only captured them accurately and agreed with ESMM for high-quality samples. We also noticed that across all replicates of these luminal epithelial cell samples, the replicability for nine out of ten states was higher for ESMM (see Additional Figure S6B).

Overall we conclude that the integration of DNA methylation in ESMM allows to robustly capture the cross-talk between different layers of epigenetic marks and enhances the definition of modeled states in previously non-defined regions.

### ESMM uncovers cell-type-specific regulatory transitions during B-cell differentiation

The role of complex epigenetic changes during the B-cell differentiation process has been a matter of intensive studies [20, 31]. Based on various data (including Hi-C based 3-D data) a model was proposed (Figure 4A) which summarizes the stepwise epigenomic transition from naive B-cells (NBC) to germinal center B-cells (GCBC) branching into memory B-cells (MBC), and plasma cells (PC) [20]. We speculated that by combining our ESMM based segmentation with matching Hi-C based segmentation data, we should get deeper insights into global and local epigenomic changes accompanying B-cell maturation.

**Figure 4:**
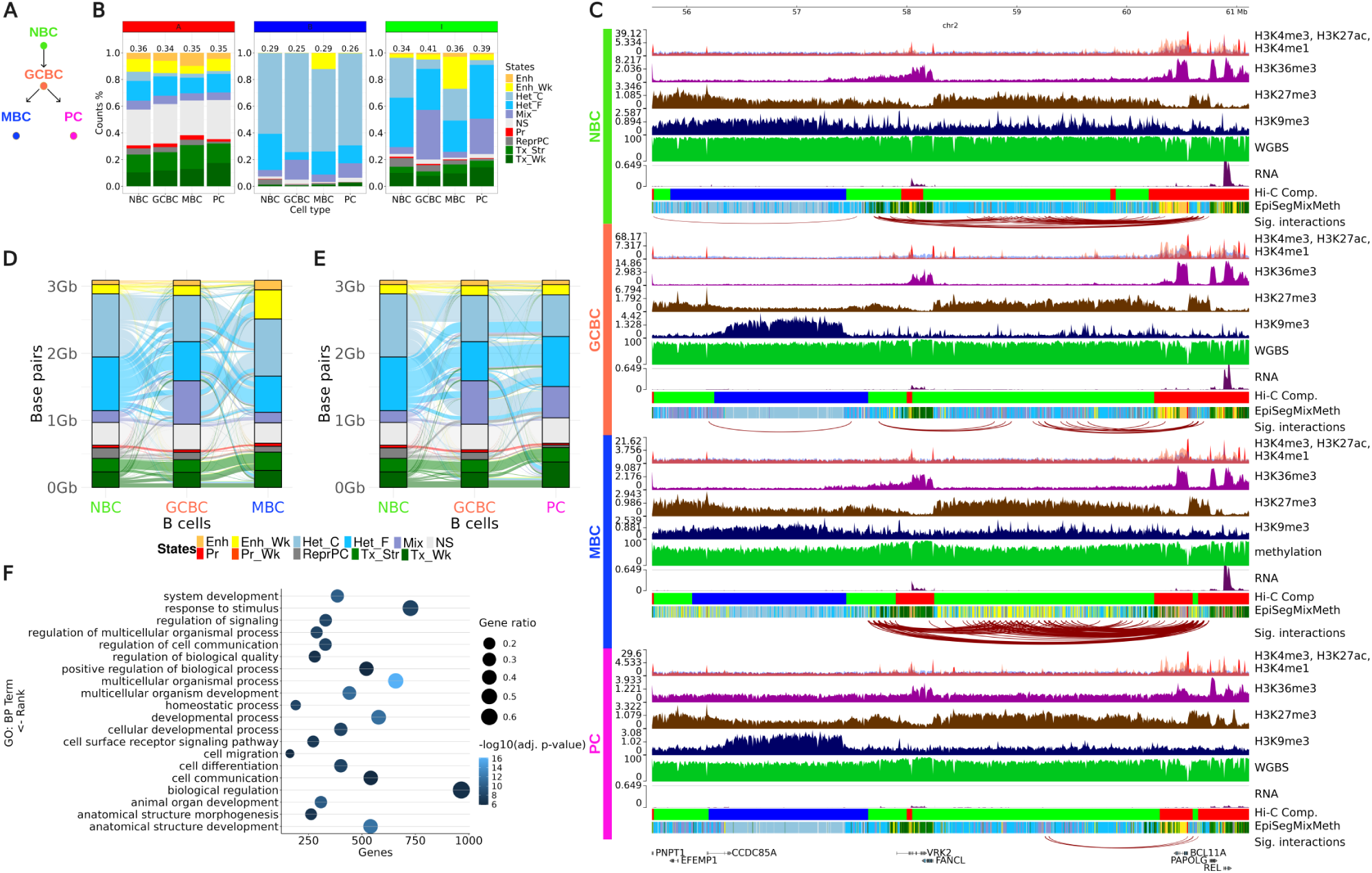
Analysis of state transtions during B-cell differentiation. (A) Current model of B-cell differentiation. (B) Distribution and proportional changes of EpiSegMixMeth states within 3D compartments (Active ”A”: red, Inactive ”B”: blue, Intermediate ”I”: green) during differentiation. (C) Maps of segmental and epigenomic changes during B-cell differentiation across B-cells in the Vrk2/Rel region: Ordering from top to bottom for each cell type: core histone modifications (narrow marks H3K4me3, H3K27ac and H3K4me1 overlayed into a single track), WGBS methylation data (0-100%), RNA-Seq expression levels (log1p normalized), 3D chromatin state segmentation, EpiSegMixMeth segmentation. The arcs below show local significant (max 1MB distance) interactions based on Hi-C contact map (thickness signifying importance). (D) Sankey diagram depicting the state transitions in EpiSegMixMeth from NBC to GCBC to MBC, with the y-axis showing genomic coverage and proportional transitions between states. (E) The state transition diagram from NBC through GCBC to PC, similar to (D). (F) Gene Ontology analysis for genes affected by facultative heterochromatin to weak enhancer transition in MBC, selected based on EpiRegio database (see Methods).

To assess how ESMM-based segmentation aligns with 3D genome architecture, we compared it to Hi-C–derived A, B, and I compartments in the same B-cell differentiation states (Figure 4B). The A and B compartments displayed relatively consistent genome coverage across cell types, ranging from 34–36% for A and 25–29% for B, whereas the I compartment showed broader variability (34–41%), reflecting its dynamic nature during B-cell maturation. Consistent with earlier findings ([20, 31]), the I compartment emerged as the most variable and responsive to differentiation-related changes. Notably, this compartment was enriched in ESMM-defined chromatin states associated with regulatory activity and heterochromatin remodeling: weak enhancers (Enh Wk), constitutive heterochromatin (Het C), and facultative heterochromatin (Het F) in MBC; and mixed states (Mix) along with Het F in GCBC and PC. These patterns indicate that the I compartment serves as a hotspot for chromatin remodeling, with ESMM providing a finer-grained and functionally informative view complementing the coarse Hi-C compartmentalization view. A deeper examination of specific genomic regions further highlights the added resolution of ESMM. In MBC, ESMM identified numerous small segments classified as weak enhancers (Enh Wk) within ”I” compartment. One such region, encompassing the Rel (NF-*κβ*) and VRK2 genes-key regulators in memory B-cell function-exhibited a transition from facultative heterochromatin (Het F) in NBC to a mixture of different states in GCBC to dispersed weak enhancers in MBC, back to a mixture of different states in PC. These finer-grained ESMM segments corresponded with increased and localized MBC-specific 3D contacts (Figure 4 and Figure S10), suggesting the formation of novel regulatory interactions and loops (depicted by arcs in Figure 4C) linking enhancers to nearby regulatory elements. In addition to enhancer resolution, ESMM revealed broader domain-level chromatin changes. For instance, an expansion of the A compartment near the VRK2 locus was observed in NBC and MBC but not in GCBC or PC. This shift was accompanied by changes in adjacent heterochromatic regions, including an increase in H3K9me3 signal and a transition from Het F to Het C in GCBC and PC. The segmentation patterns suggest lineage-specific regulatory architecture: MBC retains greater similarity to NBC, while PC more closely resembles GCBC in this region. Globally, MBC showed an increased abundance of weak enhancer states compared to NBC, GCBC, and PC (Figure 4B), and genes linked to these enhancers were enriched in functional categories related to development, signaling, response to stimuli and cell differentiation (Figure 4F).

Finally, a direct comparison of segmental changes from GCBC to MBC versus GCBC to PC (Figures 4D–E) highlights divergent regulatory trajectories. The GCBC-to-MBC transition is marked by an increase in enhancer-like segments and a reduction in the Mix state, consistent with regulatory activation. In contrast, the GCBC-to-PC transition shows an expansion of Het F, with no notable gain in enhancer activity, suggesting a shift toward chromatin compaction and transcriptional silencing in PCs. Such segmental changes are also reflected in local changes in the 3D contact maps (see Supplementary Figure S10) and are also observed on the genome-wide level between B-cell types as clear cell type specific inter- and intra-segmental contact patterns (Figure S8).

Overall our analysis shows that ESMM segmentation has the potential to capture a broad spectrum of active and repressive states, both in the long range chromatin context as well as in a local gene specific context and allows to more precisely link epigenomic segmentation to the 3-dimensional domain and interdomain interactions in human cells.

## Discussion

### Implications and advantages of combined modeling of epigenomic marks for chromatin segmentation

In this work, we summarize the results of the first integrated and comprehensive SAGA analysis across a diverse set of 154 IHEC-grade epigenomes, utilizing both histone modification and DNA methylation signals. While existing segmentation tools such as ChromHMM, Segway, and EpiCSeg have advanced the analysis of histone modifications, they typically neglect or decouple DNA methylation, limiting the biological insight that can be gained from truly integrated analyses.

To address this, we developed ESMM, an extension of our earlier method EpiSegMix (ESM) [16], which combines a flexible probabilistic modeling framework with state duration modeling and distributional versatility. ESMM incorporates both ChIP-seq–derived histone modification data and WGBS–derived methylation data, without resorting to binarization thresholds that discard quantitative variation. This is a key advantage, especially in low-methylated regions (LMRs) and heterochromatic domains, where DNA methylation carries important cell-type-specific regulatory information as shown previously [27].

Previous efforts at integrative segmentation have either relied on indirect post hoc combination of methylation and histone data or, in rare cases, simplistic binarization strategies that obscure biologically meaningful variability. For instance, one prior study in mice [32] applied a 50% methylation cutoff per 200 bp window. Such approaches are particularly problematic in regions of partially methylated domains (PMDs) or facultative heterochromatin, where methylation levels vary continuously and have fuzzy patterns. In addition, binarization impairs the comparison of segmentation results across (many) cells, as overall methylation levels vary significantly between certain cell types and disease states [5]. ESMM overcomes these limitations by modeling raw methylation counts directly, enabling quantitative integration of both signal types. Combining the resulting segmentation of ESMM with a random forest classifier enabled us to perform the first comprehensive comparative analysis of 154 primary human cell and tissue samples, which are archived as high-quality complete epigenomes of healthy cells in the IHEC EpiAtlas [4]. This comparative segmentation represents a novel and unique resource to visualize and track cell- and tissue-specific epigenomic features.

### DNA methylation enhances resolution and stability of epigenome segmentation

Conventional histone modification-based segmentation tools classify a substantial portion of the genome (up to ~ 70%) as ”no signal/quiescent” state [33, 34] leaving a large portion of the genome poorly characterized and unexplored.

Among the regions previously labeled as quiescent, ESMM reassigns 60% to heterochromatin, 30% to euchromatin, and only 10% remain as true no-signal regions, demonstrating an enhanced resolution across genomic domains.

For example, we observed that integrating DNA methylation not only facilitated a more accurate capturing of constitutive and facultative heterochromatin but also provided a deeper understanding of their cell-type-specific attributes (Figure S5). For many cell- or tissue-specific genes, we find interesting combinations of cell-specific DNA methylation signatures (PMDs and LMRs) and heterochromatic or euchromatic marks. Regions with constitutive heterochromatin, such as those controlling diverse types of allele-specific epigenetic control in all cells, including X-chromosome inactivation, allelic exclusion, and genomic imprinting [35], do not exhibit this variability across cell types and form stable and uniform PMDs.

Importantly, ESMM maintains strong concordance with histone-only segmentation like ESM and ChromHMM, but demonstrates increased stability when ChIP-seq data are noisy or incomplete (see Figure S7, jaccard index, states% and meth%). This is particularly true for broad marks such as H3K27me3 and H3K9me3, where ESMM segmentation remains reliable even when these two marks were absent, in contrast to histone-only segmentation tool. In such cases, DNA methylation was sufficient to recapitulate key heterochromatin states with high reproducibility, reflecting the strong correlation between DNA methylation and repressive histone modifications [36, 37, 38]. Moreover, ESMM properly captures the small regulatory elements like promoters and enhancers (Figure S7, jaccard index, states% and meth%).

Collectively, these findings highlight ESMM as a powerful tool for comprehensive chromatin state annotation, especially in datasets where histone marks are missing or limited but WGBS data are available.

### ESMM captures global and local regulatory epigenomic transitions during B-cell differentiation

Building on the annotations provided by ESMM, we used this for a focused analysis of B-cell epigenomes. The role of complex epigenetic changes during differentation of human B-cells has been studied quite intensively before ([20, 31]). Together with 3-D data a model was proposed (Figure 4A) which describes the stepwise transition from naive B-cells (NBC) to germinal center B-cells (GCBC), memory B-cells (MBC), and plasma cells (PC). We speculated that leveraging the existing comprehensive dataset including full epigenomes as well as matching 3D chromatin interaction data might give us a deeper insights in the epigenomic changes accompanying the differentiation. In particular, we aimed to deeper understand the changes of A, B and I compartments in differentiating B-cells based on HiC-data.

While regions marked by constitutive heterochromatin, promoters, and transcribed states largely maintain stable states (with slight shifts in borders), notable transitions occur in regions marked by facultative heterochromatin and *mixed* state. These dynamic regions often transitioned from weak heterochromatic states to weak enhancers, and in some cases reverted back to a repressive configuration, highlighting the plasticity of regulatory domains during lineage commitment. Notably, many of these transitions were only detectable through the integration of DNA methylation into the segmentation model. For example, in the Rel (NF-*κβ*), Rel-DT/VRK2 locus, we observed a marked emergence of discrete weak enhancer elements in MBCs. While ChromHMM broadly annotated this region as a single weak enhancer domain, ESMM was able to resolve it into multiple smaller elements, each associated with distinct methylation and chromatin features. This finer resolution provided by ESMM better captures the functional modularity of regulatory elements during B-cell maturation. ESMM-identified weak enhancers co-localize with an increased number of MBC-specific chromatin loops, suggesting they are functionally linked to promoter regions of flanking genes. This spatial association further supports the regulatory relevance of these newly resolved enhancer elements.

Hi-C data revealed that regions previously annotated as ”quiescent”-using histone-only segmentation models-can be more accurately reclassified into biologically relevant states using ESMM. As illustrated in Figure S8, these regions exhibit extensive long-range interactions with various active chromatin states, indicating that biologically meaningful features may have been overlooked under the earlier ”quiescent” classification.

Overall, our work demonstrates that ESMM segmentation aligns very well with previous broader segmentations, such as the A/B/I classification from Hi-C data [20] and chromatin-based segmentations. it delivers superior granularity that reveals fine-scale epigenetic transitions often masked in coarser models. This includes gene-specific regulatory contact changes, such as those observed in the SKI gene, a well-known transcriptional regulator in memory B-cell differentiation [39, 40]. In summary, our analyses demonstrate that ESMM generates data offering a multiscale view on the genome by defining the most comprehensive genome wide long range regional segmentation while also capturing local cell-type-specific regulatory elements and transitions at single-locus resolution as seen for the differentiating B-cells.

## Conclusion

The use of a flexible read count-based integrated segmentation approach offers an extended high resolution view into cell specific adaptations on a genome wide level. By incorporating both histone modification and WGBS DNA methylation data, this method enables the reclassification of up to 70% of the human genome that was previously annotated as quiescent or no-signal by other SAGA approaches. The integration of DNA methylation not only enhances the robustness and stability of chromatin state annotations, but also effectively compensates for the absence of key heterochromatic marks. The power of this integrative approach is particularly evident in its ability to resolve fine-scale chromatin changes—such as those observed during B-cell differentiation—when aligned with 3D genome architecture data, revealing dynamic transitions that would otherwise remain obscured.

### Limitations

Our approach relies on epigenomic data of sufficient quality, good coverage, and uniform processing as provided by the epiATLAS data. Particularly the coverage of DNA methylation data should be sufficient (i.e., ≥ 10x) to capture the variable distribution of CpG positions across the genome and to quantify local DNA methylation changes with high enough precision. The use of a smoothed window based approach (200pb windows) for DNA methylation annotation as applied in ESMM might therefore still lead to an underestimate of local contributions of DNA methylation variation particularly in regulatory segments.

### Outlook

ESMM is inherently flexible and can be extended to integrate additional count-based epigenomic data. Incorporating data types such as open chromatin assays (e.g., ATAC-seq, NOME-seq, or DNaseI-seq), or direct sequencing approaches that simultaneously capture genomic and DNA methylation information, may further enhance model performance. These additional layers of information can provide a more comprehensive view of the epigenomic landscape and enable deeper insights into how cells respond to developmental signals or disease-related changes, particularly in the context of underlying genomic variation.

## Data avalaibility

EpiSegMixMeth is available as part of EpiSegMix and can be downloaded from https://gitlab.com/rahmannlab/ episegmix. EpiSegMixMeth segmentation tracks will be provided as part of IHEC EpiATLAS data portal: https://ihec-epigenomes.org/epiatlas/data/.

## Author contributions

N.A. and J.S. contributed equally to this work. J.W. and A.S. conceived the study. N.A., J.W. and A.S. designed the study. The method, computational data analysis was coordinated/developed by N.A. and J.S. under the supervision of S.R., J.W., and A.S.. L.L. processed and performed the primary analysis of 3D genomic data. N.A., J.S and J.W. contributed to writing, and all authors contributed to revising the manuscript.

## Ethics declarations

None declared

## Methods

### Input data and processing

We downloaded all epigenomic data from the IHEC repository, specifically focusing on entries in the Epigenome Reference Registry (EpiRR) marked as complete epigenomes. These datasets include the six key histone marks (ChIP-Seq data): H3K4me3, H3K27ac, H3K4me1, H3K36me3, H3K27me3, and H3K9me3, along with DNA methylation data (WGBS) and gene expression signals (RNA-Seq). We selected only primary cells designated as healthy, resulting in 154 EpiRR entries that met these criteria. We acquired ChIP-Seq BAM and bigwig files, WGBS CpG.bed and bigwig files, and RNA-Seq gene.results and bigwig files from these entries. All data was aligned to the reference human genome (hg38). Using the ESM package [16], we generated the ChIP-Seq count matrices by applying a sliding non-overlapping window of 200 bp to the BAM files. For the WGBS data, we used the BSmooth method [41] to smoothen methylated reads per CpG site using a 200 bp sliding window. We then calculated the averages of methylated and unmethylated reads across CpGs for each bin to obtain bin counts (assigning 0 methylation values in the matrix if the 200bp bin has no CpG) for a total of 15 441 313 bins in the ChIP-Seq count matrix. We included counts of chromatin marks and the average methylated and unmethylated read counts for each bin in the final count matrix. For ChromHMM, we binarized the input signals using the BinarizeBam command from the ChromHMM package [42], with the 200 bp window size parameter and hg38 as the reference human genome.

### Hi-C data preprocessing, normalization, and interaction calling

Hi-C data was obtained from the European Genome-Phenome Archive [43] (accession number EGAS00001004763). The sequencing reads of Hi-C experiments were processed with the nf-core/hic pipeline, version 2.0.0 [44], *genome* set to ‘GRCh38’, *digestion* to ‘mboi’, *resolution* and *binsize* to 1000, 2000, 5000, 10000, 20000, 25000, 30000, 35000, 40000, 45000, 50000, respectively. Read quality control was performed by FastQC [45]. The main

HiC-Pro module of the pipeline performed the mapping (using bowtie2 with a two step strategy to rescue reads spanning the ligation sites), detection of valid interaction products, duplicate removal and generation of raw and normalized contact maps. Genome-wide contact maps were created at different resolutions, using cooler v0.10.2 [46]. The cooler balancing algorithm was applied to normalize the contact maps [46]. Biological replicates (rep1, rep2 and rep3) were merged with the cooler merge option and default parameters.

### Compartment and TAD calling

The data was binned using *genome binnify* at the aforementioned resolutions from cooltools v0.7.1 [47]. Afterwards, the matrix was corrected for GC bias and very low/high contact regions, with default parameters for *genome gc* from cooltools. Compartments were called with cooltools *call-compartments*. The compartments were called against intra-chromosomal data only instead of genome-wide, with the flag --cis-only; otherwise, default parameters were used. TAD calling was performed with hicFindTADs from the HiCExplorer tool suite v3.7.2 [48, 49, 50], with parameters --minDepth 300000 --correctForMultipleTesting fdr.

### Log2-ratios of normalized interactions

Normalized Hi-C maps were analyzed at different resolutions, from the four B-cell subpopulations. Logarithmic ratios of the normalized contact map data were computed between NBC and GCBC, between GCBC and PC and between GCBC and MBC, with the hicCompareMatrices tool from HiCExplorer [48, 49, 50].

### EpiSegMixMeth, EpiSegMix and ChromHMM segmentation

We used binarized signals of core histone marks from the ChromHMM BinarizeBam command as an input for the LearnModel command from the ChromHMM package to perform ChromHMM segmentation with parameters of binsize 200 bp, numstates 10, and assembly hg38.

As an extended version of ESM, ESMM allows each signal to be fitted independently with a different type of probability distribution in the mixture model. ESMM supports fitting Poisson, Binomial, Negative Binomial, Beta Binomial, Beta Negative Binomial and Sichel distributions to model histone counts [16]. We fitted the best distribution type for each chromatin and DNA methylation dataset. To consider both coverage and methylation information, ESMM fits DNA methylation count data using either the Binomial or Beta Binomial distribution, where the number of trials corresponds to methylation coverage and the number of successes to the number of methylated CpGs. The input for EpiSegMix and EpiSegMixMeth are the histone count matrix and DNA methylation level per 200 bp genomic bin (see Input data and processing). We ran EpiSegMix and EpiSegMixMeth with duration modeling (online available workflowTopology/Snakefile) using default parameters and setting states to 10. We used the Beta Negative Binomial (BNB) distribution to fit the read counts of all six histone marks. In EpiSegMixMeth, we used the Beta Binomial (BB) distribution to fit DNA methylation data. We selected the Beta Negative Binomial and Beta Binomial distribution due to their high flexibility and good performance in previous experiments.

We trained 154 individual models representing each sample. For the reduced model (EpiSegMixMeth, EpiSegMix and ChromHMM), we performed the same steps, excluding H3K9me3 and H3K27me3 from the input count matrix for each model, and used only B- and T-cell cell epigenomes (40 samples).

### Annotating segmentation states with relevant functional labels

For each sample, we constructed an enrichment overlap incorporating 23 features derived from three models: EpiSegMixMeth, EpiSegMix, and ChromHMM. The analyzed features included mean overlap counts for histone modifications H3K4me3, H3K27ac, H3K4me1, H3K36me3, H3K27me3, and H3K9me3; methylation ratio; genomic coverage of each state; CpG density; mean gene expression; and overlaps with genes, exons, transcription start sites, transcription end sites, TSS 2kb upstream, PMDs, and various families of repeat elements such as SINE, RNA, Retropson, LINE, LTR, RC, and Tandem repeats. We normalized these features using a min-max approach for each sample to calculate an enrichment score ranging from 0 to 1. Utilizing these scores, we manually assigned functional labels to each of the 10 states across 29 samples, each representing a different cell type. We employed a random forest classifier [51] in R (4.3.1), training it on these 290 data points with the ‘ntree’ parameter set to 100. To guarantee unique labeling within each sample, we utilized the Linear Sum Assignment Problem’s [52] (LSAP) cost matrix. For states assigned with low probability, we designated exclusive labels for the sample within the models.

### Finding state overlaps and epilogos

To calculate the state overlap between EpiSegMixMeth and ChromHMM’s *no signal* state, we counted each 200 bp bin marked as the ChromHMM *no signal* state across any of the 154 samples. We then overlaid the ChromHMM-ns annotated bedfile with each EpiSegMixMeth state for all samples and calculated the mean overlap counts for each state across all samples. To assess the importance of states in EpiSegMixMeth and ChromHMM using saliency features based on information entropy, we utilized Epilogos[53] with the single mode (–single) and saliency level 1 (S1 - Kullback-Leibler relative entropy). We subsequently visualized the output from Epilogos using pyGenomeTracks[54] around the AICDA locus (chr12:8,550,000-9,650,000).

### Predicting gene expression using RNA-Seq

To assess the predictive power of gene expression based on segmentation labels from each model, we overlapped the gene.results file for each sample, assigning each bin its log-transformed gene activity (FPKM). We then split the data into training and testing sets in a 70-30% ratio, respectively, and trained a linear model to predict gene activity using labels as categorical variables. Finally, we quantified and reported the model’s accuracy using the coefficient of determination (R^2^) and Root Mean Square Error (RMSE) of the test set.

### Complete vs reduced model analysis

To measure the similarity between states defined by the complete and reduced models, we employed the Jaccard index from bedtools [55] as a similarity score. For each state in the reduced model, we assigned the corresponding state in the complete model that had the highest Jaccard index, ensuring that each label was used only once per sample. Additionally, to generate data for Sankey plots, we counted the combinations of state pairs from the reduced and complete models for all 200 bp bins across the dataset.

### Consistent annotation within same samples of heterogeneous quality

Consistent annotation within the same samples of heterogeneous quality We analyzed eight samples of luminal epithelial cells, which varied in quality based on the H3K27me3 metric. We categorized these samples into three groups: low, mid, and high quality. As we used samples derived from the same cell type, we treated them as biological replicates and expected them to have consistent segmentation annotations, despite differences in quality. For each EpiRR, we calculated the proportion of all bins marked as facultative heterochromatin and plotted this proportion against the quality of H3K27me3, measured by Jenson-Shannon distance, in increasing order. Furthermore, we fit a smooth linear regression line between the genomic coverage of facultative heterochromatin and the quality metric of H3K27me3, and plot this regression line along with the variance for both the EpiSegMixMeth and EpiSegMix models. To assess the consistency of annotation in samples of luminal epithelial cells, we calculated the Jaccard index score for each model (ESMM and ESM) between the same state for each pair of samples. Furthermore, we calculated the difference in similarity scores for each state between the models as JI*_ESMM_* − JI*_ESM_*. A difference *>* 0.2 indicates that ESMM provides consistent annotations, while an absolute difference of ≤ 0.2 suggests that both ESMM and ESM perform comparably well.

### Inter and intra state interactions

We overlapped Hi-C data with our state annotation to understand the interactions of states in a 3D space. For each B-cell type, we used a 200 bp resolution of normalized Hi-C data. We considered a contact valid if a 200 bp bin has a contact to another 200 bp bin in the normalized Hi-C sparse matrix (.cool files), and the linear distance between these bins is not greater than 4 Mb (megabases). We then annotated each valid contact to and from a 200 bp bin using the segmentation annotations generated by both EpiSegMixMeth and ChromHMM.

### Gene ontology using transition using ESMM defined states transistion

We overlapped the bins annotated as Het F in NBC as well as GCBC with Enh Wk in MBC. We used the coordinates of these overlapped bins in the EpiRegio database to find the genes associated or affected by these regions. Using the list of genes, we performed Gene Ontology analysis using gprofiler ([56]) and filtered only terms related to biological processes (BP). We further sorted them using the combined rank of log10(adjusted-p-value) and precision (proportion of genes for the terms).

## Abbreviations

ChIP-Seq: chromatin immunoprecipitation followed by sequencing
ESM: EpiSegMix
ESMM: EpiSegMixMeth
GCBC: germinal center B-cells
JSD: Jenson-Shannon Distance
LMRs: lowly methylated regions
MBC: memory B-cells
NBC: naive B-cells
PC: plasma cells
PMDs: partially methylated domains
RNA-Seq: RNA sequencing
TES: transciption end site
TSS: transcription start site
UMRs: unmethylated regions
WGBS: whole genome bisulfite sequencing

## Acknowledgments

We thank the International Human Epigenome Consortium (IHEC) for providing access to reprocessed and harmonized epigenomic data from a broad collection of human cell types.

## Supporting Information

**Figure S1:**
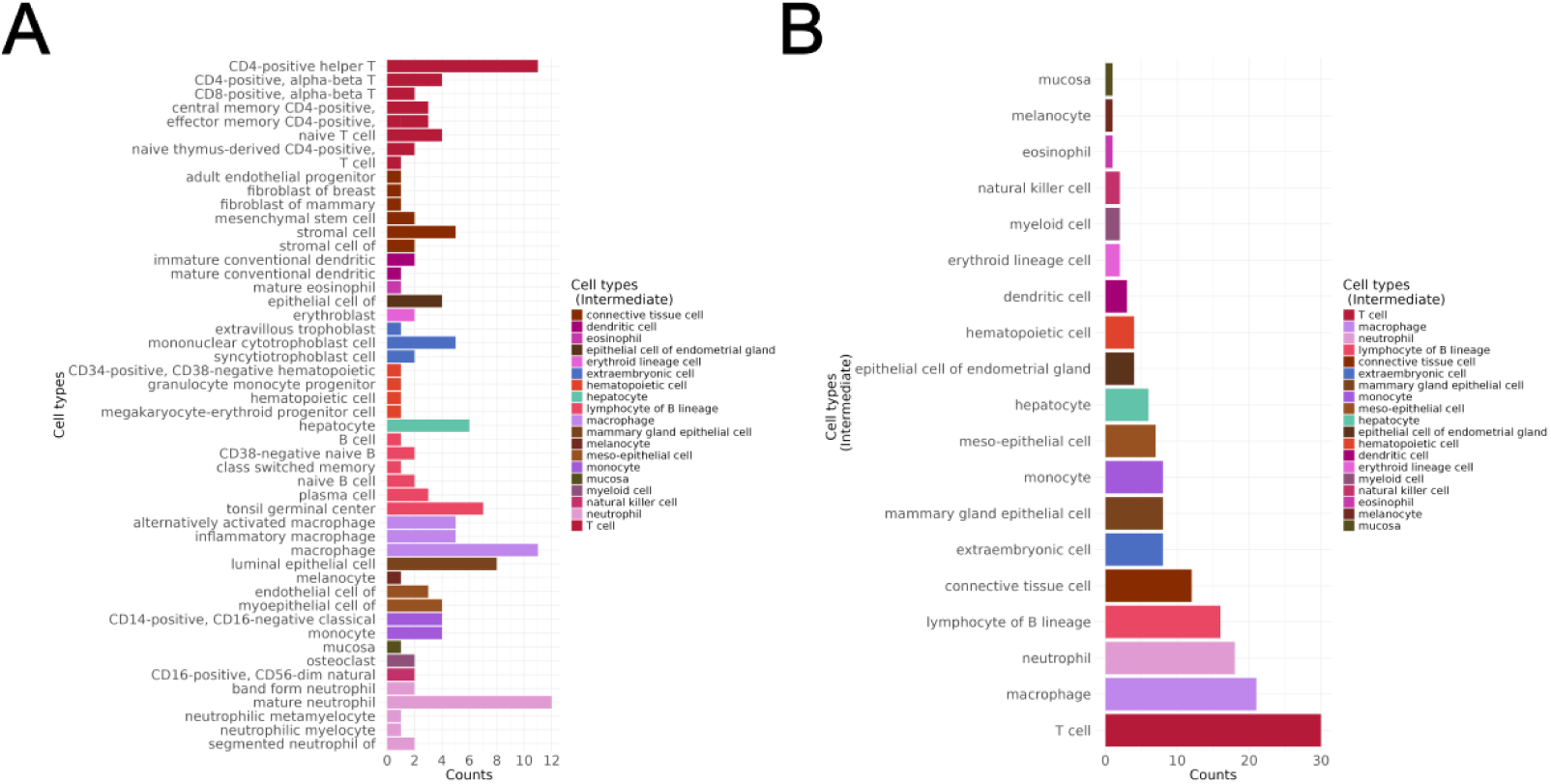
Classification of cell/tissue types across the 154 full epigenome samples of this study. (A) 50 different cell types with lower classification based on IHEC annotation, x axis showing number of samples present for each cell type annotation from various consortia merged in IHEC. (B) IHEC intermediate annotation of the 154 samples into 19 cell/tissue groups used in this study.

**Figure S2:**
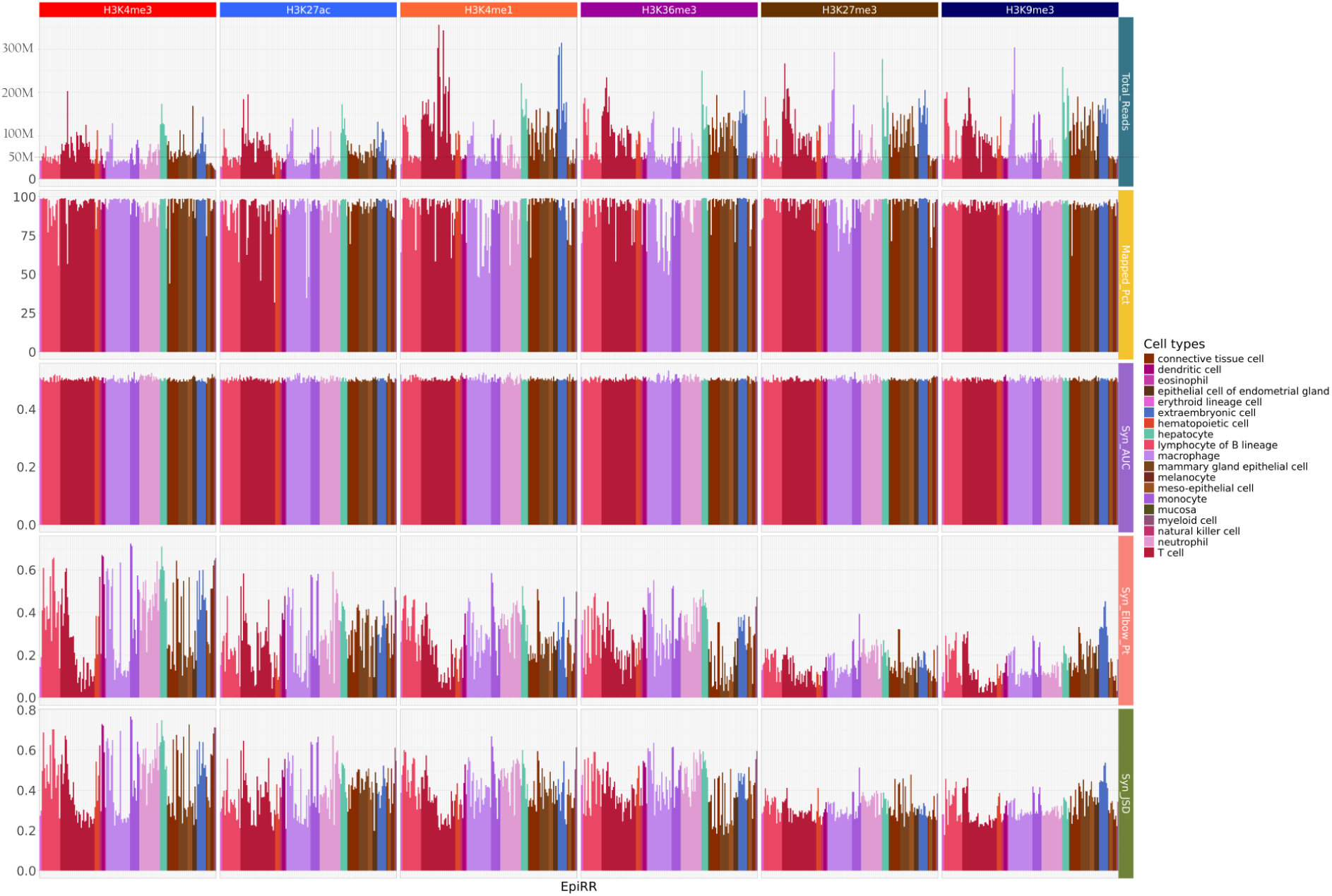
ChIP-Seq QC: Five Quality control metrics for six core histone marks are shown for 19 cell types used in the study. Top to bottom: Total reads in millions, Mapped reads percentage from total reads, synthetic Area Under Curve with max value 0.5, synthetic elbow point with max one and min 0, and synthetic Jenson-Shannon Distance with min 0 and max 1.

**Figure S3:**
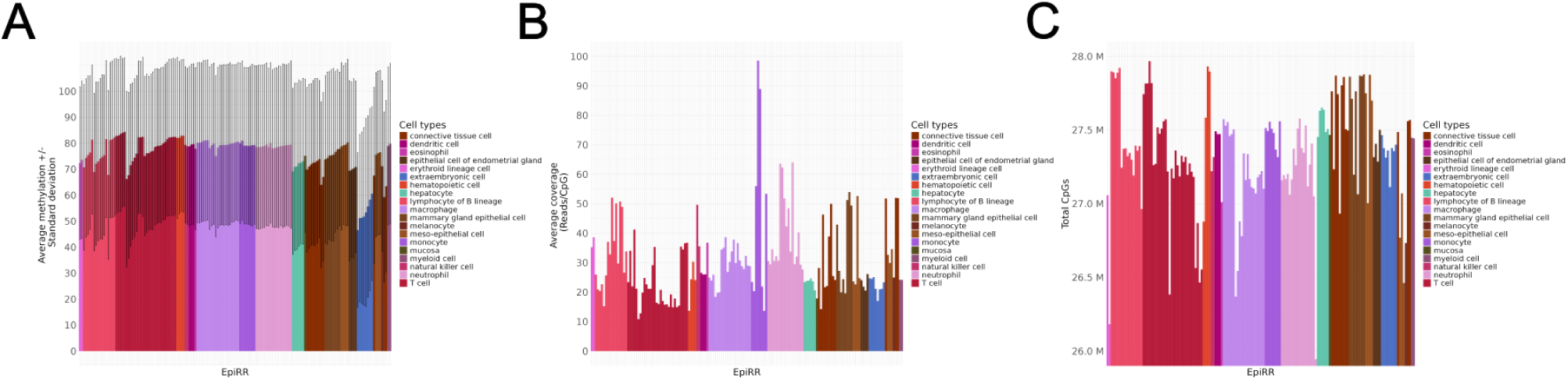
WGBS QC: (A) Average DNA methylation (0–100%) sorted by cell types and increasing average methylation within cell types across 154 samples, capturing cell-type-specific methylation signatures across 19 distinct cells types. (B) Total CpGs in million covered by WGBS for each sample, categorized by cell type and sorted by increasing average methylation levels, with a minimum of 26 million CpGs in a neutrophil sample. (C) Average sequencing depth/coverage for WGBS data measuring average number of reads covered by CpGs with sample in the same order as in (A) and (B), reaching up to 100x coverage for monocycle samples.

**Figure S4:**
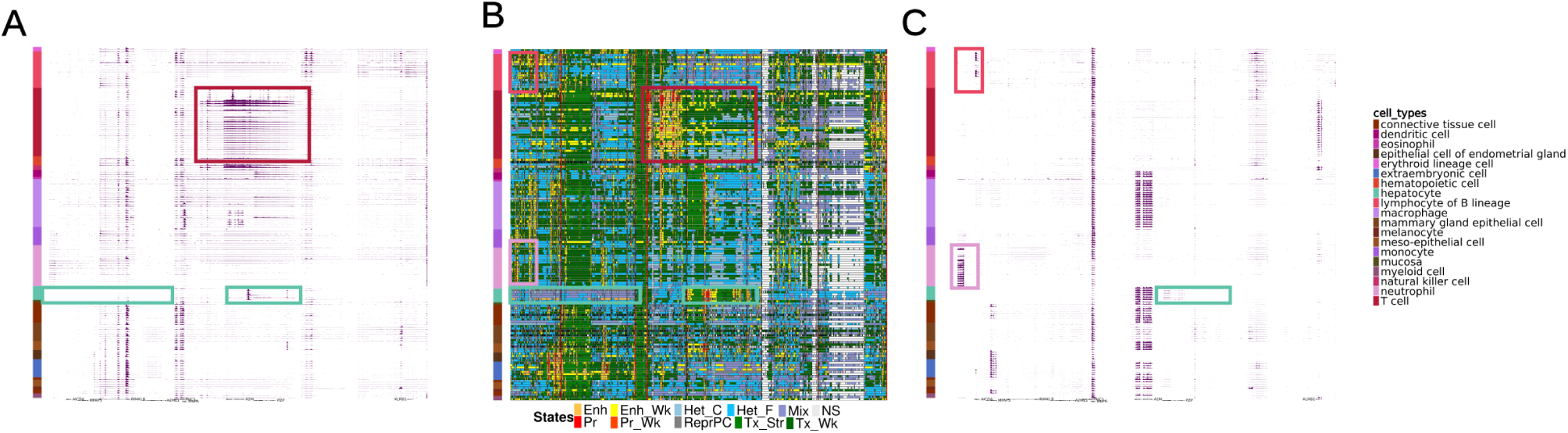
Cell-type specific segmentation of ESMM in comparison to matching RNA-Seq patterns: (A) Gene expression intensity on the reverse (minus) strand across all 19 cell types. The red box demarcates the activity of the A2M gene in a subset of immune cells, the turquoise box marks cell-specific repression of various genes in hepatocytes. (B) EpiSegMixMeth segmentation tracks in the same order as (A), highlighting the boxed overlay of epigenetic states with RNA-Seq signal. Note the strong expression in (A) accompanied by the accumulation of regulatory states (yellow and red segments). Conversely the absence of gene activity in hepatocytes (A) with heterochromatin states (blue and purple). The pink box indicates the B-cell-specific activity of the region surrounding the AICDA locus, as depicted in (C). (C) Same as (A) but capturing gene expression activity transcribed in the forward strand, with boxes showing cell-type specific pattern corresponding to (B).

**Figure S5:**
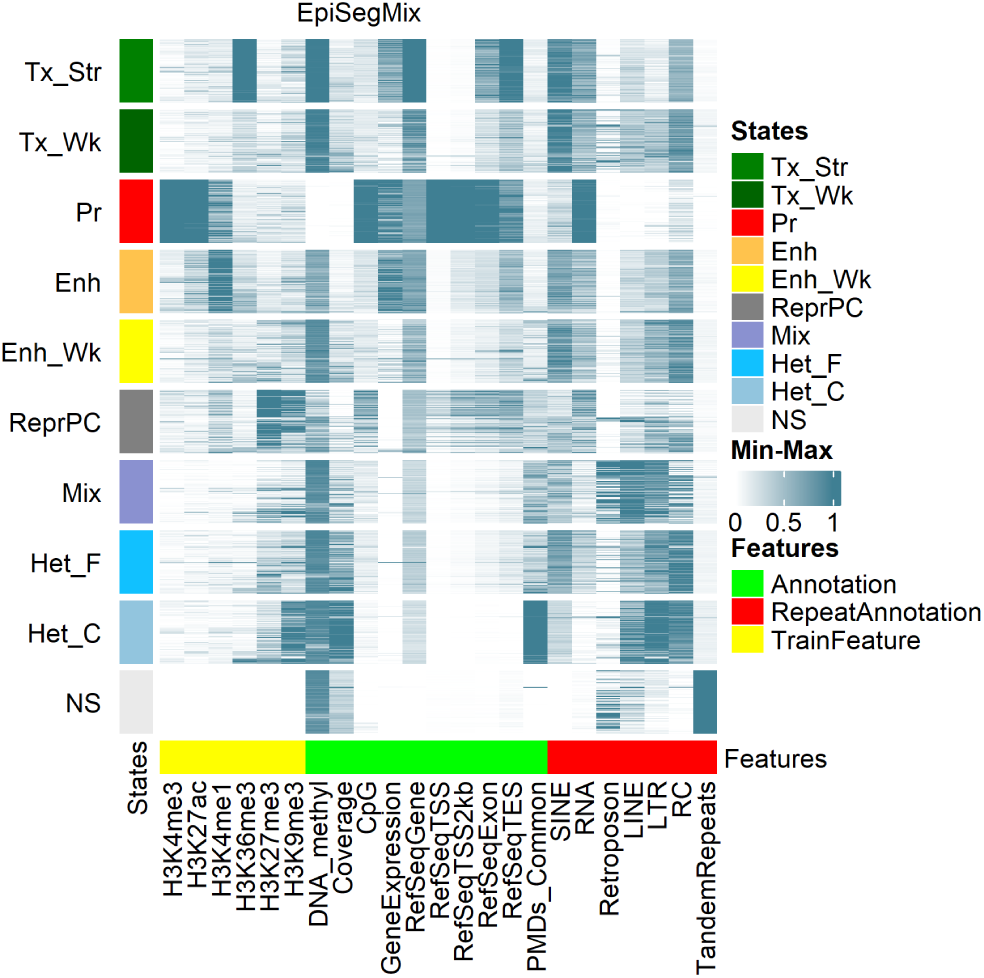
EpiSegMix states defined using six core histone marks annotated with biological functions. Labels are assigned based on the random forest classifier. Features on the x-axis are grouped into train (yellow), which are used in the model; annotation (green); and repeat annotation (red).

**Figure S6:**
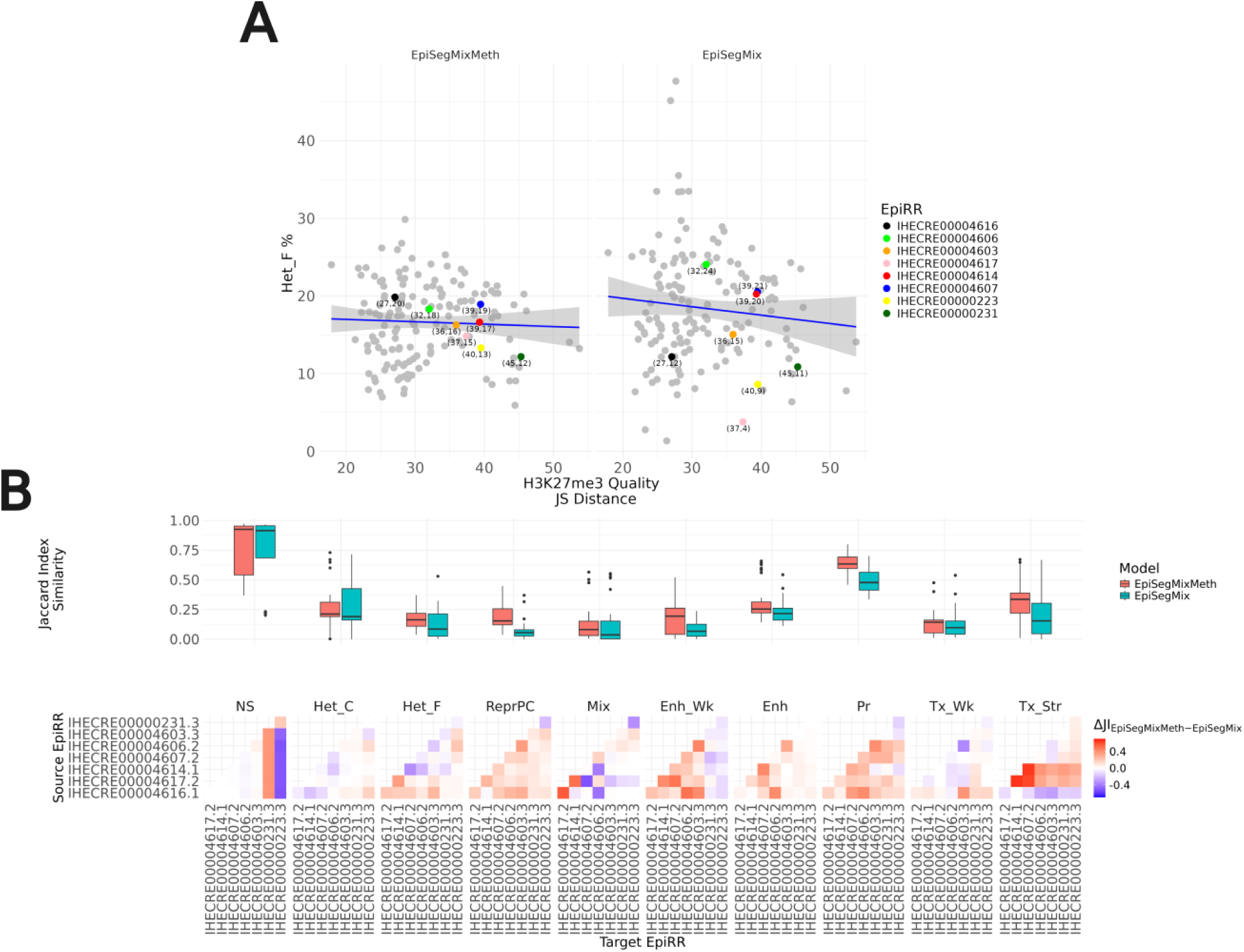
DNA methylation stabilizes segmentation calling in samples with reduced H3K27me3 ChIP signal quality : (A) *Left* : Eight samples from the same cell type, luminal epithelial, showing the quality of histone mark H3K27me3 on the x-axis, measured by synthetic Jenson-Shannon distance with samples ordered in increasing quality of H3K27me3 and genomic coverage of facultative heterochromatin (Het F) defined by EpiSegMixMeth ESMM on the y-axis as expected effect state by H3K27me3. The smooth linear regression fit between the Het F state coverage and quality of H3K27me3, shown in blue, with variance. *Right* : Same as *Left*, for states defined using EpiSegMix ESM. (B) *Up*: Boxplot comparing the ESMM and ESM models for all luminal cell samples per state (x-axis), assessing robustness using the Jaccard index (y-axis). High scores indicate that the genomic regions annotated are consistent across samples for that model. *Bottom*: Heatmap comparing EpiSegMixMeth (ESMM) and EpiSegMix (ESM) performance per sample across all 10 epigenetic states. Red/blue denotes relative ESMM out- or under-performance, respectively, against ESM by annotating consistency of regions (Jaccard Index); white indicates comparable performance.

**Figure S7:**
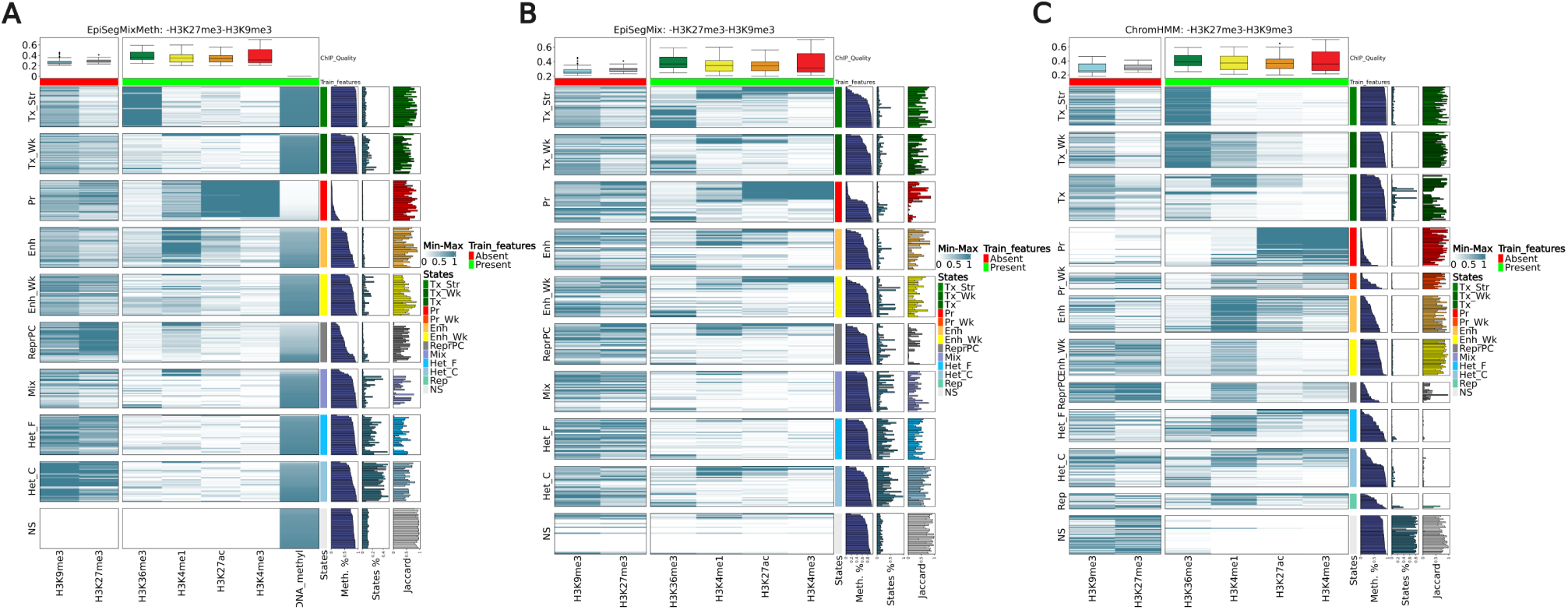
ESMM with DNA methylation (A) supports a range of regulatory and broad state recoveries epigenomes lacking H3K27me3 and H3K9me3. The figure shows a comparison to chromatin only based models (B), EpiSegMix and (C) ChromHMM. Composition of each subfigure: Heatmap displaying the enrichment of present histone marks (green bar) plus DNAmethylation (only for ESMM) and absent heterochromatic histone marks (red bar). Each row represents one of the 40 T- or B-cell epigenome samples, and rows are grouped into states to enhance state-specific enrichments. Emissions within each state are ordered in relation to the increasing averaged DNA methylation per state. Box plots above the heatmaps display the synthetic Jenson-Shannon distance of histone marks across these samples as quality metrices. On the right to emission profiles we show the following Annotations. Left: Average methylation per state for each sample. Middle: Genomic coverage of the state within the sample. Right: Similarity score (Jaccard index) measured between the state in the reduced model and the corresponding state in the complete model defined by core histone marks and DNA methylation for each sample.

**Figure S8:**
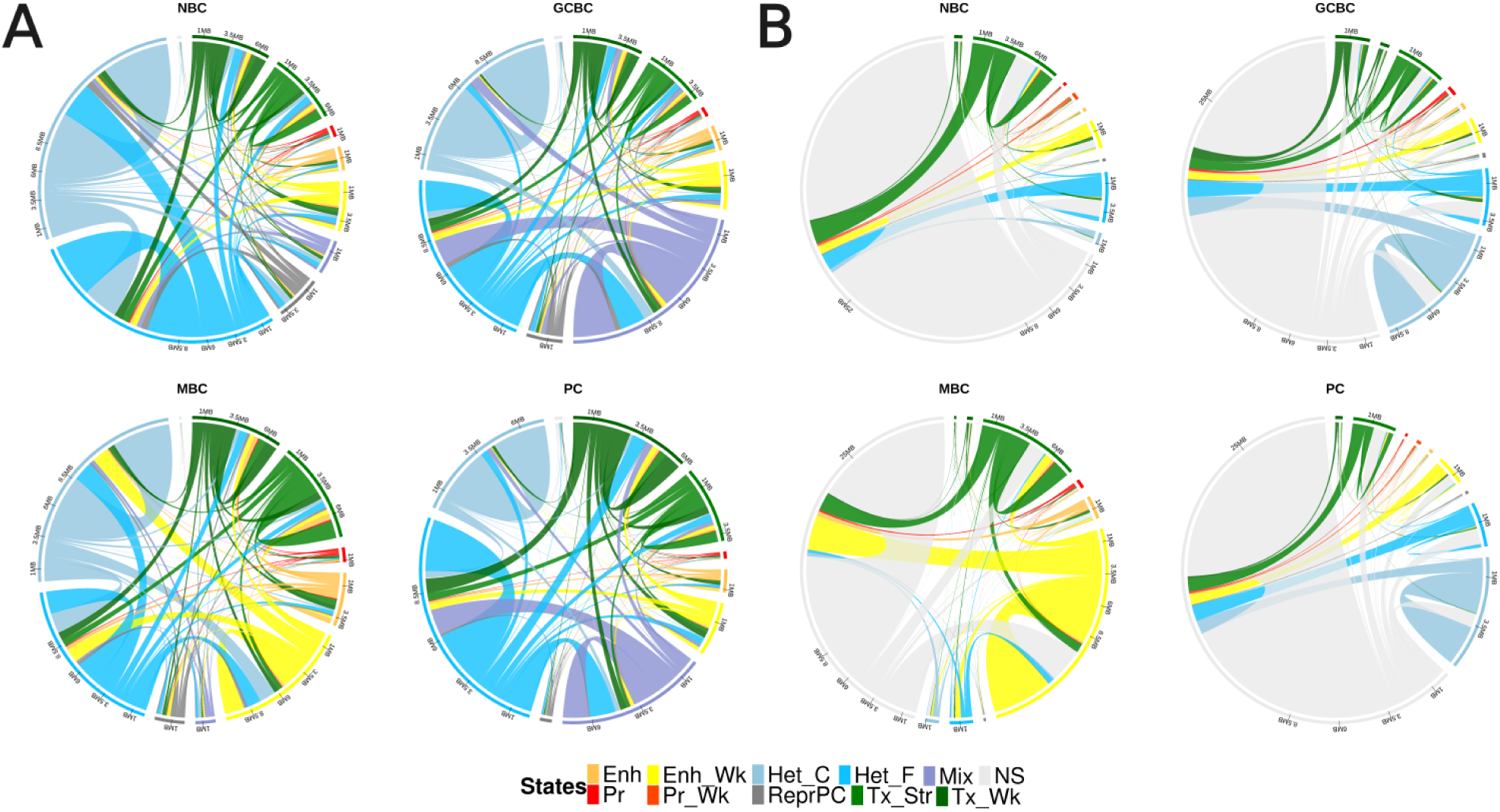
3D contacts in various B-cells in relation to ESMM(A) and ChromHMM (B) segmentation: (A) The colouring highlights the Inter- and intra-state 3D contacts in relation to ESMM segmentation. Contact are derived from normalized Hi-C data with a max distance of 4 MB between positions. The cell specific Circos plots summarise the contacts between and within states, for naive B-cells (NBC), germinal center B-cells (GCBC), memory B-cells (MBC), and plasma cells (PC). (B) Similar to (A), for states defined by ChromHMM.

**Figure S9:**
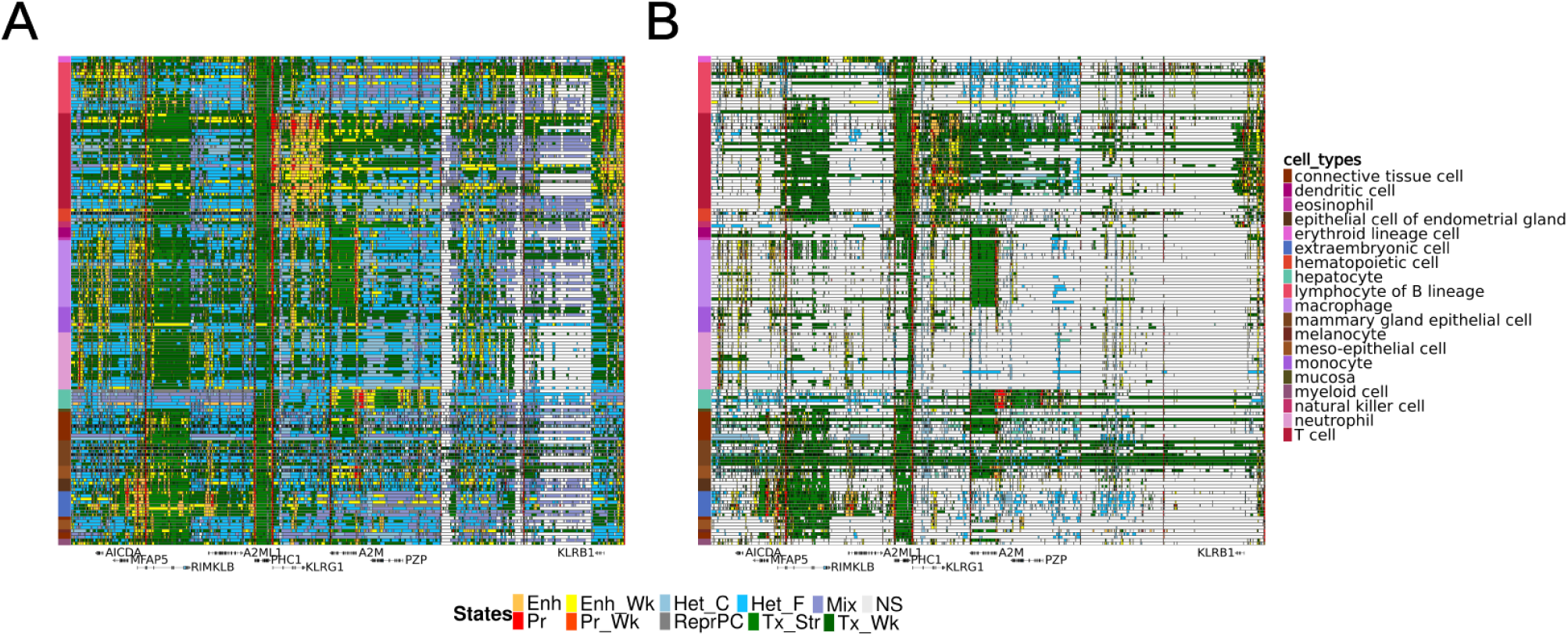
Comparison of ESMM and ChromHMM cell-type specific ordered segmentation patterns: (A) EpiSegMixMeth segmentation tracks as shown in Fig.1 D (chr12:8550000-9650000) (B) ChromHMM segmentation tracks across this region grouped in the same order as (A). Note the extensive non-signal and Het-C and Het-F states in (B) particularly for monocyte and macrophage samples.

**Figure S10:**
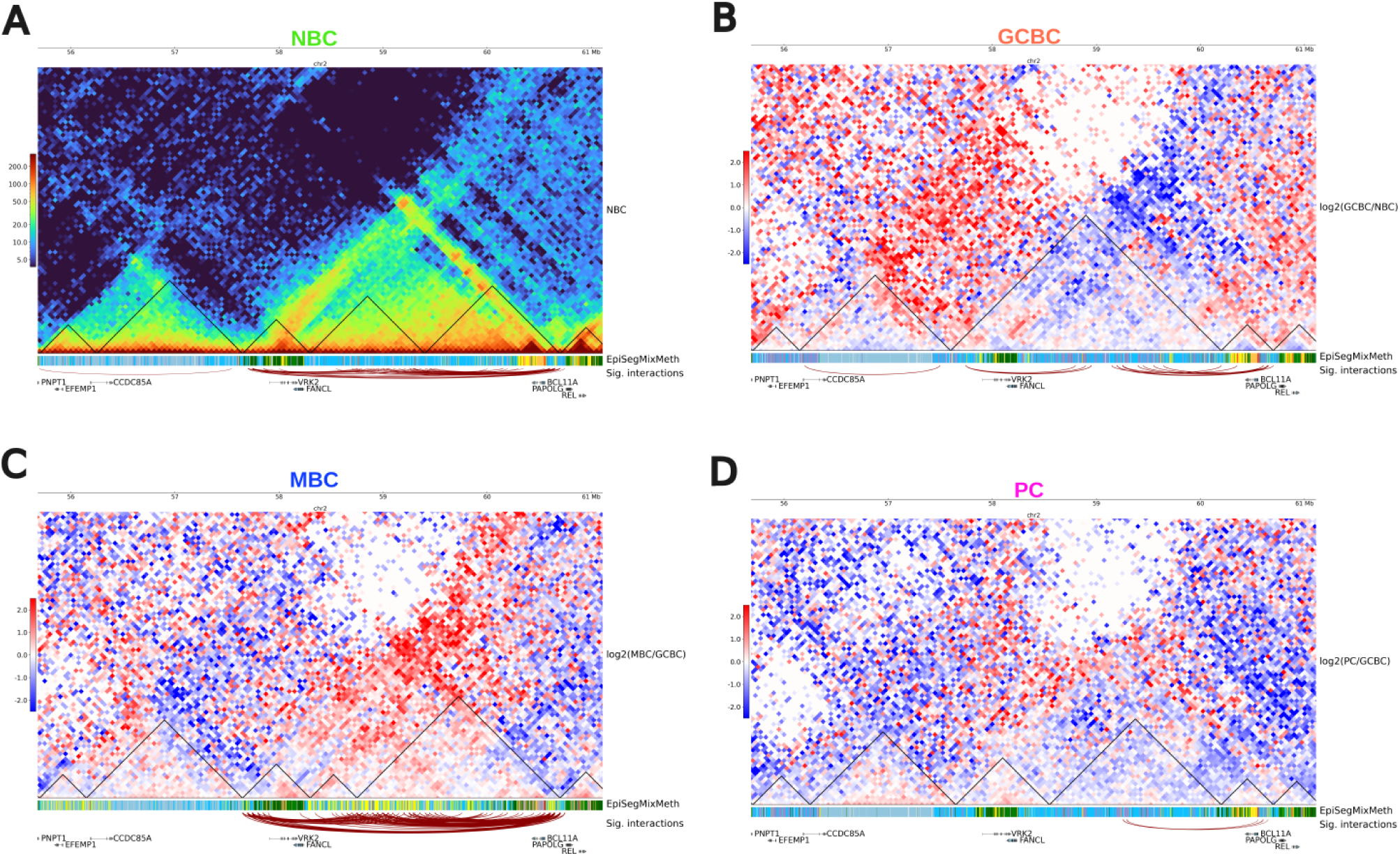
B-cell Differentiation states transition (A) Sequentially from top to bottom: Hi-C contact maps of NBC with TAD calls (black triangles), EpiSegMixMeth segmentation, and significant 1MB interactions (thichness signifies importance). (B) Similar display as (A) for GCBC, except Hi-C maps show log fold changes from Naive B-cells, with red indicating increased interactions and blue indicating reductions. (C, D) As per (B) for MBC and PC, respectively, changes are relative to GCBC.(J) The circular plot for the SKI gene displays significant interactions as arcs colored by cell type, with arc height indicating significance. Additional tracks from outermost to innermost include genomic coordinates, cell-type-specific colors, SKI gene expression levels, and exon (orange) overlaid with EpiSegMixMeth states for promoters (red), enhancers (yellow), and facultative heterochromatin (light blue), illustrating their role in gene expression changes through notable interactions.

